# Defining Hospital Catchment Areas Using Multiscale Community Detection: A Case Study for Planned Orthopaedic Care in England

**DOI:** 10.1101/619692

**Authors:** Jonathan M Clarke, Mauricio Barahona, Ara W Darzi

**Author notes:** Corresponding Author: Dr Jonathan M Clarke, Centre for Health Policy, 10^th^ Floor QEQM Building, St Mary’s Hospital, Praed Street, London W2 1NY, United Kingdom.

## Abstract

**Background:** The English National Health Service 5-year Forward View emphasises the importance of integration of hospital and community services. Understanding the population a hospital serves is critical to formulating strategies for community engagement and determining their accountability for populations. Existing methods to define catchment areas are unable to adapt to dilute health care markets in urban areas where populations may interact with several different hospitals. Formulating catchment areas which permit the inclusion of more than one hospital based upon patient behaviour allows for collaboration between hospitals to reach out into the communities they collectively share.

**Method:** The proportion of presentations from all census Middle Super Output Areas (MSOAs) to every hospital trust providing orthopaedic care in England were calculated. The cosine similarity of all MSOAs to one another was computed from these proportions. Multiscale community detection was applied to planned orthopaedic surgical admissions in England from 1st April 2011 to 31st March 2015. Stable community configurations were identified and the proportion of patients presenting to hospitals located within the catchment area in which they resided was calculated. The performance of these catchment areas was compared to conventional methods for assigning mutually exclusive catchment areas.

**Results:** 2,602,066 planned orthopaedic surgical admissions were identified for patients resident in 6,791 MSOAs in England attending 140 different hospital trusts. Markov multiscale community detection revealed five stable catchment area configurations consisting of 127, 51, 26, 15 and 11 catchment areas. Between 78% (127 catchments) and 93% (11 catchments) of clinical presentations were to hospitals within a patient’s allocated catchment area compared to 76% for the “first past the post” method.

**Conclusions:** Multiscale community detection is a novel and effective, data-driven method for defining mutually exclusive, collectively exhaustive catchment areas in secondary care. In urban areas with dilute healthcare markets, the model favours collaboration between hospitals in covering a clearly delineated but shared catchment, and thereby produces simplified and more representative catchment areas.

## Introduction

For a health system to effectively provide comprehensive clinical care to its patients it is essential to define the relationships between hospitals and the communities of patients they serve. These relationships are generally referred to as ‘catchment areas,’ and the different ways in which they are defined has drawn significant interest from the fields of economics, geography, health policy and statistics over recent decades.^1–5^ A catchment area represents an association between the provider of a service and its users. In the healthcare setting, users may represent individual patients, or groups of patients, identified by shared clinical, demographic or geographic traits (most commonly geographic aggregation at the level of the zip code or census division).

The definition of catchment areas is composed of two stages: association and partition. Geographic units and care providers are quantitatively associated to one another by means of spatial proximity or patterns of clinical presentation.^6,7^ Such associations between users and hospitals constitute the ingredients to be evaluated, and either retained or removed, in order to define the structure and partitions into catchment areas. The principal decision in the partitioning lies in whether catchment areas should be mutually exclusive and collectively exhaustive (MECE), or, alternatively, based on *a priori* thresholds of association (e.g., distance from a hospital, frequency of usage, etc).

Creating a set of catchment areas that are mutually exclusive (i.e., without overlap between catchments) results in a partition that is straightforward to represent geographically, and which represents an unambiguous link between hospitals and communities of patients.^1^ If associations derived from clinical presentation data are used, the hospital attended most frequently by a group of users defines the allocation to MECE catchment areas. This method, known as ‘First Past the Post’ (FPTP), thus assigns a geographic area to the provider to which it sends the largest number of patients.^4,6,8^

An alternative approach to defining catchment areas is to identify all groups of users for which the strength of association to a hospital exceeds a defined threshold. For example, the catchment area for a hospital may be all areas where more than a defined percentage of clinical presentations from within that area are to that hospital. Alternatively, from the perspective of the hospital, its catchment area may be defined as all geographic areas that contribute more than a defined percentage of their total case volume.^3,9^ In contrast to MECE methods, there is no guarantee that such criteria will allocate all geographic regions to a catchment area, and, additionally, catchment areas may overlap one another.^1^

The literature to date lacks a dominant method of determining the catchment areas of health care providers, partly resulting from the many purposes for which catchment areas are used. Some applications require the partition of a region or country into MECE regions, while other require a more comprehensive understanding of the communities from which demand is drawn, accepting the overlap of catchment areas as a consequence. Such contradicting requirements pose a challenge for different methods. For example, in urban areas, where there is a high spatial density of hospitals, and patients from the same geographic areas interact with more than one hospital, the conventional paradigm of assigning one hospital to one catchment area presents various shortcomings.^10^ Mutually exclusive methods will fail to capture any clinical attendances aside from the most commonly attended hospital, whereas threshold-based partitions produce either highly overlapping catchment areas, in which hospitals have indistinct accountability for care provision to a community, or stringent catchments that leave large geographic regions unallocated.

In the following sections, we introduce a method to produce mutually exclusive, collectively exhaustive catchment areas, which circumvents some of the limitations of existing approaches. The method is based on Markov Stability multiscale community detection, which uses Louvain optimisation to detect and obtain robust partitions that are intrinsic to the network, without imposing an *a priori* level of coarseness.^11–15^ The approach takes a dynamical viewpoint to detect such relevant partitions; it analyses how a Markov process spreads on the network over (Markov) time and establishes the presence of communities (groups of nodes) within which information is shared quickly and retained over long times. Such a signature signals a robust partition, which can be found at different levels of coarseness (larger and larger communities of nodes) as the Markov time increases.. These techniques have been used to identify communities within processes as diverse as metabolic networks, transport systems and power grids.^13–15^ Here, we use this methodology to extract intrinsically robust partitions at different levels of resolution from the network of MSOA regions of England (nodes) linked to one another by weighted edges reflecting not any geographic information but the similarity of their patterns of hospital utilisation (see Methods).

While it is naturally assumed that the catchment areas of hospitals will consist of multiple geographic subunits, it is also assumed that a catchment area must only contain a single hospital. As health services become more centralised and the overlap between traditional catchment areas increases, it is appropriate that catchment areas consisting of multiple hospitals may be realised. Indeed, in densely populated urban areas, several hospitals offering similar services exist in close proximity to one another. In such circumstances, First-Past-the-Post methods would exclude many patients who live in areas where a particular hospital is not its most frequently attended hospital. Conversely, thresholded methods would be able to capture the areas from which a hospital’s cases are drawn, but would produce catchment areas that are so overlapping with one another as to be of limited interpretation.^1^ As a result, catchment areas in such regions are either not representative or not useful.

The multiscale community detection process overcomes this difficulty by identifying regions that correspond to dilute markets and combining catchment areas in such a way as to produce mutually exclusive, yet representative catchment areas consisting of more than one hospital where there is significantly overlapping patterns of hospital utilisation.

## Methods

This study aims to apply multiscale community detection to planned orthopaedic surgical admissions data from England to produce mutually exclusive, collectively exhaustive catchment areas which permit collaboration of hospitals in providing coverage for catchment areas. Planned orthopaedic surgery occurs at a high volume across England and is provided by a majority of acute hospital trusts. Its incidence is such that over the study period, all Middle Super Output Areas are expected to contribute patients. It has been used as a model for the investigation of other modelling processes across health policy and health economics.^1,17^

### Data Preparation

All elective admissions under the specialty of orthopaedic surgery for adults resident in England to hospitals located in England between 1st April 2011 and 31st March 2015 were identified from Hospital Episode Statistics obtained from NHS Digital. Providers were identified at the level of the hospital trust, and where organisations merged or separated during the study period, they were treated as a merged entity throughout the study period. The co-ordinates of the population weighted centroids of each MSOA were obtained from the Office for National Statistics.^18^ Co-ordinates for hospital trusts were identified directly using Google Maps. All data processing was conducted using Python (Python Software Foundation), which was used for data extraction and analysis. Matlab version 2018 (The MathWorks, Inc.) was used for the multiscale community detection process.

### Calculation of Market Concentration

The concentration of healthcare markets at the level of the MSOA was examined using the Herfindahl-Hirschman Index (HHI). This index is widely used to assess healthcare market competition in the United States of America, but is rarely used to evaluate naturally occurring markets in public healthcare systems.^19–21^ The HHI is defined as the sum of the squares of the market shares of each hospital within a market, which in this case is defined by a single MSOA, as shown in Equation 1. The HHI ranges from an upper limit of 1, indicating only a single provider within a market, to a lower limit of 1/N, where N is the number of providers active in the market.

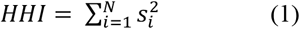

**Equation 1:** Herfindahl-Hirschman Index. Where s_i_ is the proportion of patients from an MSOA attending hospital i, and N is the set of all hospitals in the dataset.

### Generation of Undirected Unipartite Cosine Similarity Networks

For each MSOA, the proportion of admissions to each hospital was calculated, forming an M by N matrix, where M is the number of MSOAs and N is the number of hospitals. The cosine similarity of each MSOA to every other MSOA in the dataset was calculated according to Equation 2.

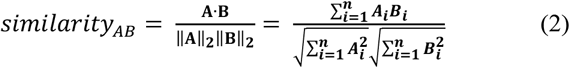

**Equation 2:** Calculation of cosine similarity between MSOAs. Where A_i_ is the proportion of presentations from MSOA A to hospital i and B_i_ is the proportion of presentations from MSOA B to hospital i.

This produces an M by M matrix which in turn may be considered as an undirected unipartite weighted network consisting of a node for each MSOA with edges connecting each MSOA weighted according to their cosine similarity to one another. This cosine similarity network was sparsened through the application of the ‘Relaxed Minimum Spanning Tree’ process to generate a network consisting of only ties that are informative to its local and global structure.^11,22^

### Multiscale Community Detection

This sparse cosine similarity matrix may then undergo Markov Stability Multiscale Community Detection to identify communities of geographic regions that are most similar to one another according to patterns of flow in the original bipartite network. Through applying multiple Markov chain processes across the network, communities of different scales may be identified across increasing Markov time.^13,14^ Through this method, communities of varying scales ranging from 6791 communities each containing a single MSOA to one community containing all MSOAs may be generated. Markov Stability Multiscale community detection was run over a range that reveals community structures of similar magnitude to the number of hospitals included in the dataset. In this case this corresponds to Markov time from 10^−2^ to 10^3^ with 500 time-steps. Each time-step was simulated 500 times. Markov times associated with stable partitions of the network are then reanalysed at 100-fold greater temporal resolution to identify the most stable partition within each stable region.

### Evaluation of Multiscale Community Detection Output

The Markov stability process used to generate communities produces three outputs for each time step. For a detailed review of each parameter and its generation see Schaub et al. (2012) and Amor et al (2015).^11,14^ The outputs are as follows:

#### Number of Communities

This value describes the number of communities that the network is partitioned into at each time step. Communities are defined as subgraphs of the larger network such that the Markov process is more concentrated within that community than would be expected by chance. Generally, the number of communities decreases monotonically over increasing Markov time.

#### Variation of Information

The Markov stability method simulates partitioning 500 times and thereby generates 500 optimised partitions for each time step. The variation of information is the average of the pairwise difference between each of the 500 optimal partitions.^23^

The Markov stability method does not seek to produce the best partition, instead presenting the characteristics of optimal partitions across time steps. Subjective interpretation of the output is therefore necessary to identify across time steps which partitions represent meaningful, stable community structures. Generally, such structures are said to exist where there is low variation of information and the number of communities formed is stable across an extended time span in Markov Time..

### Evaluation of Partitions

The proposed method produces communities of Middle Super Output Areas according to their similarity in patterns of presentation to hospitals for planned orthopaedic care. As such, the process does not enforce a relationship between a hospital and a catchment area. Instead, hospitals are assigned to the geographic catchment area within which they are spatially located. Therefore, a catchment area may potentially contain no hospitals, one hospital or several hospitals. The coverage of each catchment area (C_i_) is measured as the proportion of cases arising from each catchment area that are treated by hospitals located within the same catchment area (Equation 3).

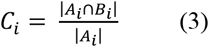

**Equation 3:** Catchment area coverage. Where A_i_ is the number of cases arising from the population within catchment area i over the study period, B_i_ is the number of cases presenting to hospitals within catchment area i over the study period. A_i_ ∩ B_i_ is the number of cases arising from the population within catchment area i that are incident on a hospital within that catchment.

The performance of each stable partition was compared to the ‘First Past the Post’ method to define catchment areas, in which each MSOA was assigned to the hospital to which it sent the highest proportion of its patients.^1^

### Regional Case Studies

Patterns of access to hospital care are known to vary according to geography and the spatial density of hospitals.^10,24^ To examine the performance of the proposed model in different geographic settings, a rural and urban case study populations were defined. The rural case study involved three contiguous counties in the South and South West of England; Oxfordshire, Gloucestershire and Wiltshire consisting of 282 MSOAs with a total population of 2.19 million people according to the most recent census in 2011.^25^ Greater London was used as an urban case study, containing 983 MSOAs with a total population in 2011 of 8.17 million people. The performance of each stable partition of the community detection model was compared to the ‘First Past the Post’ method for both regions.

This study received local ethical approval through the Imperial College Research Ethics Committee (17IC4178). As this was a national, retrospective study of routinely collected administrative data, informed consent from each human participant was not required.

### Data Availability

The hospital and MSOA constituents of each catchment area are available from the authors on request.

### Code Availability

Code written in Python version 3.6 and Matlab version 2018 is available upon request from the authors. Code is written specifically to analyse Hospital Episode Statistics in England.

## Results

### Summary Statistics

A total of 2,602,066 admissions for elective orthopaedic surgery in England between 1st April 2011 and 31st March 2015 were identified from Hospital Episode Statistics (HES). All 6,791 Middle Super Output Areas (MSOAs - mutually exclusive, collectively exhaustive geographic census divisions with a mean population of 7,800 people) in England were included in the study. A total of 140 separate hospital trusts were included. The number of admissions per MSOA ranged from 53 to 1,135 with a mean of 383 and standard deviation of 146. The number of admissions per hospital ranged from 3,791 to 50,244 with a mean of 18,586 and standard deviation of 9,468. We used the Herfindahl-Hirschman Index (HHI) defined in (1) to characterise market concentration. The mean HHI for England was 0.66 (range 0.15 to 1.00, SD 0.19), with local clusters of high HHI observed across the country (Figure 1).

**Figure 1:**
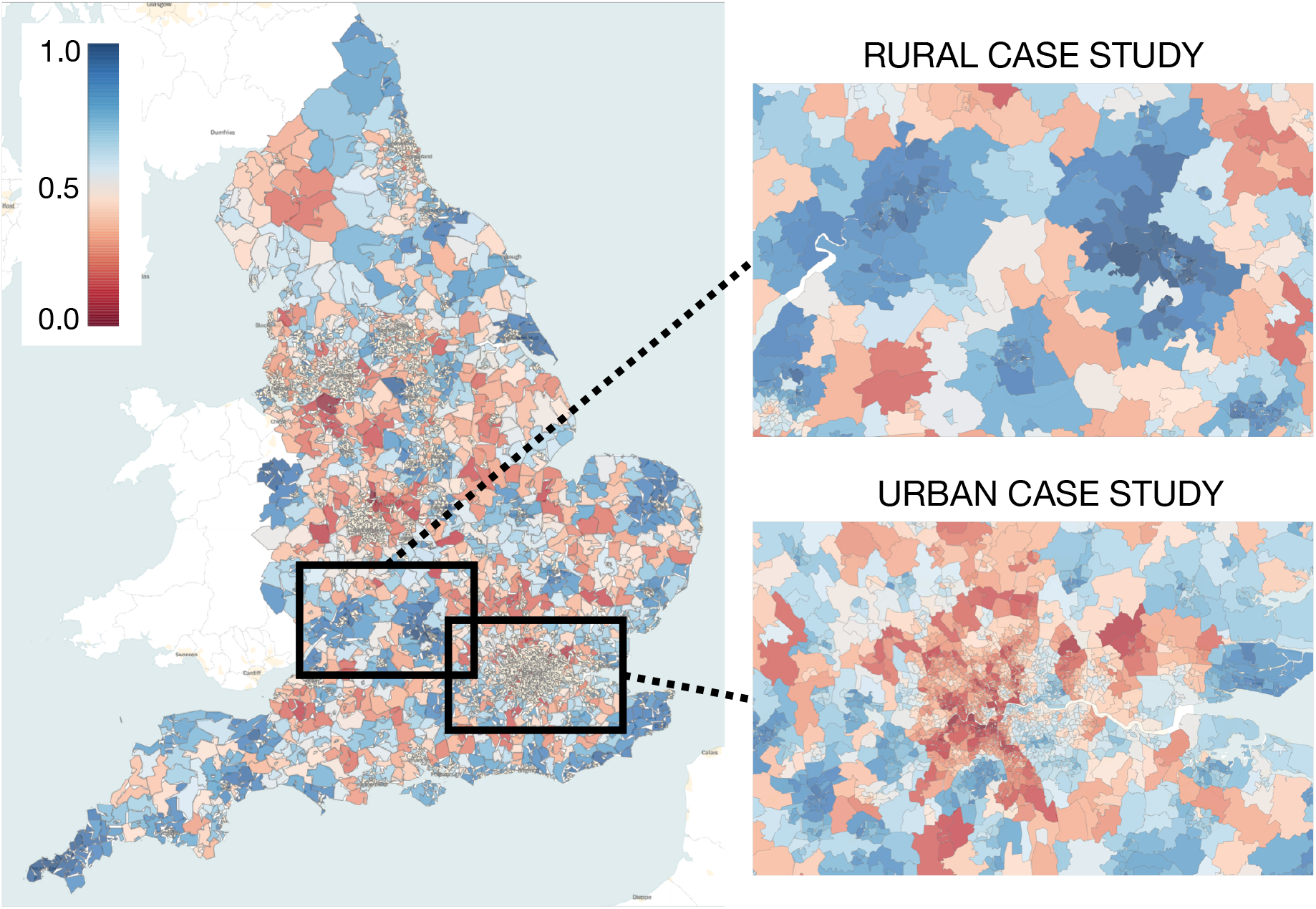
Herfindahl Hirschman Index (HHI) for elective orthopaedic care for each MSOA in England. Dark blue indicates concentrated markets, while red indicates dilute markets. The areas featured in the ‘Rural Case Study’ and ‘Urban Case Study’ are included in pop-out figures on the right-hand side.

For the rural case study, a total of 100,163 admissions for planned orthopaedic care occurred during the study period. MSOAs in this region had a mean HHI of 0.79 (range 0.30 to 1.00, SD 0.18). There are three distinct regions of high HHI, corresponding to highly concentrated markets, as shown in Figure 1. For the urban case study, a total of 281,692 admissions for planned orthopaedic care occurred. MSOAs in this region had a mean HHI of 0.48 (range 0.15 to 0.89, SD 0.16). HHI is low across the region, particularly in West London, as shown in Figure 1.

### Description of MSOA Output

Figure 1 shows the number of communities and their robustness (given by relative minima of the variation of information) as a function of increasing Markov Time. The number of communities in the partitions declines monotonically as Markov time grows, and the variation of information shows five relative minima, indicated by vertical grey bands, which underwent finer analysis at 100-fold higher temporal resolution. These five bands correspond to scales with robust partitions at different levels of coarseness. Table 1 shows the Markov time and the number of communities of the partition associated with the lowest variation of information within each of these five scales.

**Table 1:** Characteristics of the five stable partitions of the MSOA cosine similarity matrix using Markov Multiscale Community Detection.

|  | Markov<br>Time | Number of<br>Communities | Variation of<br>Information |
| --- | --- | --- | --- |
| Partition A | 0.156 | 127 | 0.01222 |
| Partition B | 3.541 | 51 | 0.00912 |
| Partition C | 21.68 | 26 | 0.00084 |
| Partition D | 157.4 | 15 | 0.00424 |
| Partition E | 464.5 | 11 | 0.00205 |

Table 2 shows the overall congruence of each partition in comparison to the First-Past-the-Post (FPTP) method. A total of 137 catchment areas were identified by the FPTP method nationally, out of 140 providers, indicating that three hospitals were not modal providers of orthopaedic care to any MSOA in England. Each of these three hospitals was a small provider of orthopaedic care located close to larger care providers. Across England, 76% of patients underwent orthopaedic surgery in the hospital contained within their FPTP catchment.

**Table 2:** Coverage and number of catchment areas in England, the rural case study and the urban case study following Markov Multiscale Community Detection in comparison to the First Past the Post method.

|  | England |  | Rural Case Study |  | Urban Case Study |  |
| --- | --- | --- | --- | --- | --- | --- |
|  | Coverage | Catchments | Coverage | Catchments | Coverage | Catchments |
| First Past the Post | 0.760 | 137 | 0.790 | 8 | 0.584 | 22 |
| Partition A | 0.775 | 127 | 0.875 | 8 | 0.630 | 21 |
| Partition B | 0.860 | 51 | 0.901 | 7 | 0.864 | 10 |
| Partition C | 0.899 | 26 | 0.924 | 5 | 0.905 | 6 |
| Partition D | 0.932 | 15 | 0.968 | 3 | 0.944 | 3 |
| Partition E | 0.934 | 11 | 0.933 | 4 | 0.962 | 2 |

Nationally, over the five stable partitions produced by Markov Multiscale Community Detection, the proportion of cases presenting to hospitals within the defined catchment areas increased from 77.5% to 93.4% as the number of catchment areas reduced from 127 to 11 (Table 2). An increase of 9 percentage points is noted between Partition A (77.5%) and Partition B (86.0%). The geographic distribution of these catchment areas for each partition is shown in Figure 3. Note that no information about the relative geographical location of the MSOAs was used to obtain the partitions with Markov Multiscale Community Detection, which are based exclusively on hospital utilisation data.

**Figure 2:**
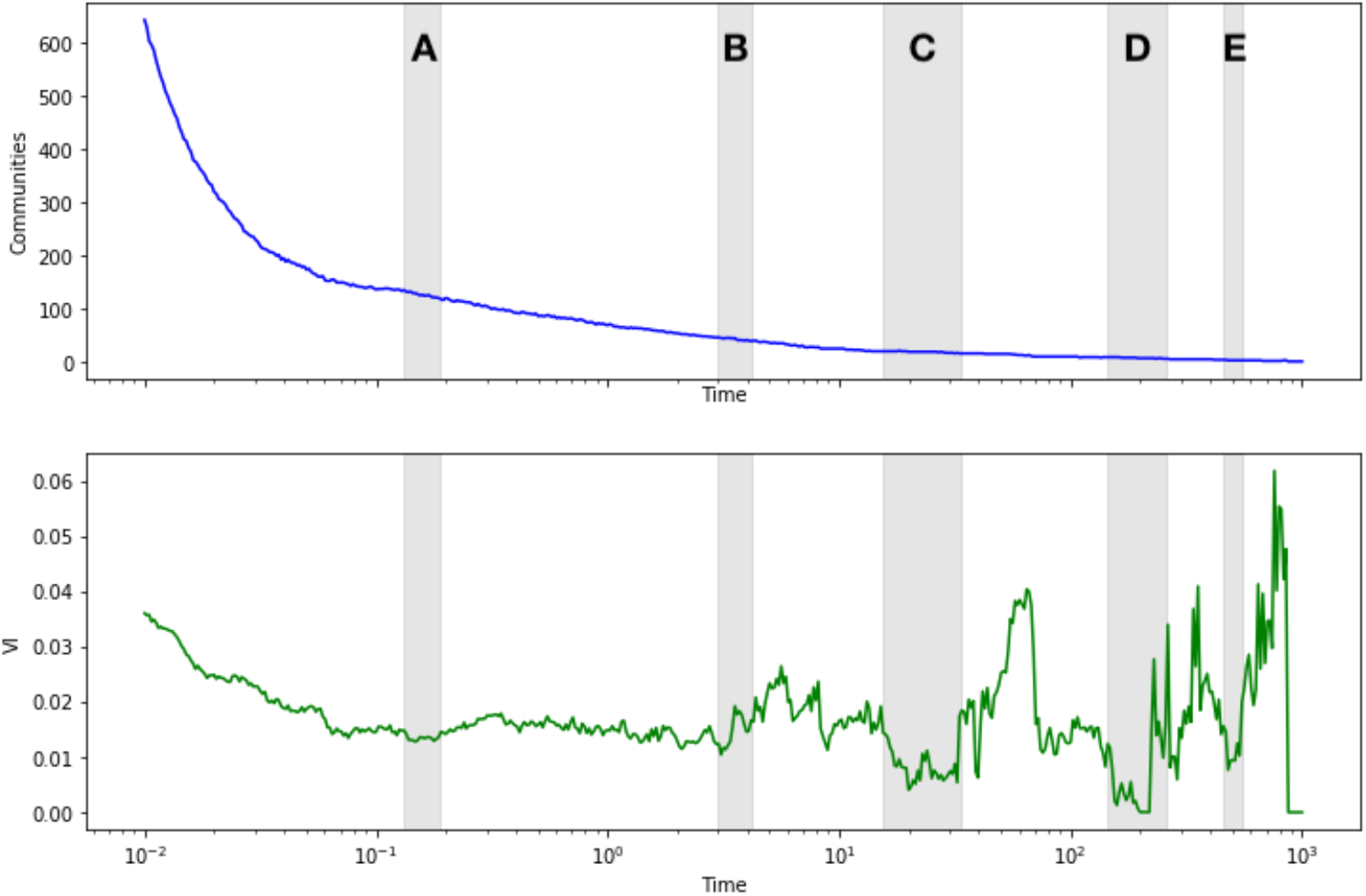
Markov Multiscale Community Detection output for planned orthopaedic care. Grey bands indicate Markov times associated with low variation of information (green line) and a stable number of partitions (blue line). These regions underwent reanalysis at 100-fold temporal resolution to determine local minima in variation of information.

**Figure 3:**
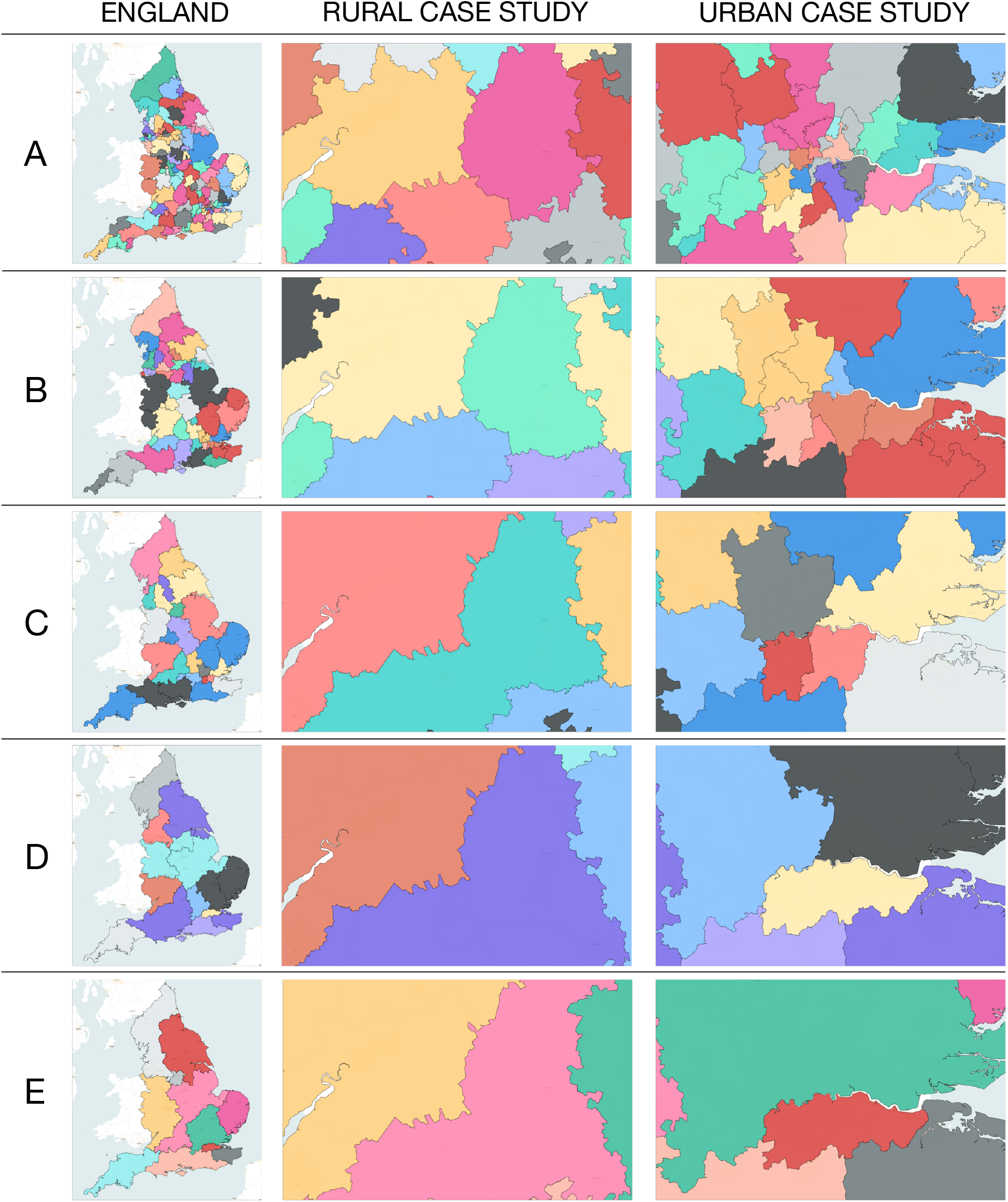
Geographic representation of the catchment areas produced from Markov Multiscale Community detection for each of the five stable partitions (A-E). The left-hand column shows the distribution of catchment areas across England, while the middle and right-hand columns show catchment area allocation for the rural and urban case studies respectively.

In the rural case study, the FPTP model produced 8 catchment areas in the region, with 79.0% of patients presenting to hospitals located within their catchment area. Over the five partitions produced, the number of catchment areas decreases from 8 to 3 before increasing to 4 in the final partition. Coverage increases from 87.5% to 96.8% before decreasing to 93.3% in the final partition.

In the urban case study, the FPTP model produced 22 catchment areas in the region, with 58.4% of patients presenting to hospitals located within their catchment area. Over the five partitions produced, the number of catchment areas decreases from 21 to 2. Coverage increases from 63.0% to 96.3% over the five partitions. An increase of 23 percentage points is noted between Partition A (63.0%) and Partition B (86.4%).

## Discussion

In the literature to date there is no dominant method of determining the catchment areas of health care providers. This is partly a consequence of the diverse applications of catchment areas and the contrasting requirements imposed on such partitions. Some applications require mutually exclusive, collectively exhaustive regions, and methods such as First Past the Post models are used to clearly identify accountable communities for a provider.^4,6,8^ Other applications of catchment areas require a more comprehensive understanding of the communities from which demand is drawn and the ensuing overlap of catchment areas. In such cases Proportional Flow models are more appropriate.^1^

Multiscale community detection methods offer an opportunity to relax the restrictions of traditional mutually exclusive catchment area assignments. Because our method scans across all levels of resolution to find partitions of intrinsic robustness, such methods allow the flexibility for catchment areas to include more than one hospital, such that mutually exclusive catchment areas can account for the dilution of markets in densely populated urban areas. This property of Multiscale Community Detection methods is especially important in urban areas, where, market dilution hampers our understanding of the relationship between hospitals and the communities they serve.^1,10^

Creating catchment areas with more than one hospital in urban areas allows for collaboration between institutions in covering a defined geographic region, and therefore a defined population. The allocation of secondary care providers to defined populations creates accountability on the part of hospitals for their communities. In doing so, hospitals may reach out more effectively into the communities they serve. In regions where catchments are shared between hospitals, they may collaborate in an organised fashion, pooling resources and sharing capacity to limit duplication and enhance the reach of secondary care interventions in the community. The increase in catchment area coverage in London from a set of 21 catchment areas (63%) to 10 catchment areas (86%) indicates the potential advantage local collaboration between care providers to achieve population coverage may achieve.

Partition A (with 127 communities as catchment areas) is the closest of our partitions to the level of coarseness of the FPTP results (137 catchment areas) and to the number of NHS trusts (140) in our dataset. To this level of resolution, Partition A produced a modest improvement of 2% coverage as compared to the FPTP method, which is achieved through the aggregation of a few of the hospitals with the most densely overlapping patient populations to collectively share larger catchment areas. Increased coverage in both the rural (11%) and urban (8%) case studies are similarly observed. Note that Partition A for both case studies has almost identical number of catchment areas to the FPTP partition (8 *vs* 8 for the rural case; 21 *vs* 22 for the urban case). Hence the improvement in coverage is the result of the adaptable clustering based on the intrinsic information contained in the hospitalisation usage data that underpins the network used in our community detection method.

As the robust Markov Stability partitions become coarser, there is a general increase in the coverage achieved. This is expected, as fewer, larger catchment areas, each containing more hospitals, would be more likely to account for the clinical presentations of patients resident within the geographic boundaries of these larger catchment areas. Notable increases in coverage are already achieved by modest coarsening of the catchment areas via aggregation of hospitals with largely overlapping patients, as obtained directly with community detection. For the England case, increases in coverage of 13% and 18% are achieved in Partitions B and C, respectively, where the ensuing catchment areas contain on average 2.7 and 5.4 NHS trusts. In the rural case study, just a minor coarsening from 8 FPTP catchment areas to 7 catchment areas defined by community detection (Partition B) increases coverage by 14%. In the urban case study, coarsening by a factor of two (from 22 FPTP to 10 community detection catchments in Partition B) increases coverage by 48%. Such local trade-offs to produce better national coverage are evident in other locations and in other partitions of this network, and reflect the intrinsic variation uncovered by multiscale community detection.

The stable community structures obtained from multiscale community detection are derived solely from the patterns of presentation of patients requiring planned orthopaedic care in England in an unsupervised manner, and are not informed by administrative boundaries, geographic proximity, or pre-existing relationships between hospitals. Yet the ability of the multiscale community detection process to produce stable conformations at different levels of granularity leading to the formation of a quasi-hierarchical community structure translates in this case to the combination of catchment areas in a consistent manner with geographical and organisational patterns as seen in Figure 3. In doing so, partitions of differing granularity may be used for different policy purposes: granular conformations resulting in more catchment areas may be used for local policy decisions regarding collaboration between individual institutions, while coarser representations may organisation of many institutions at a regional level. It is important to remark that the proposed method remains agnostic to the optimal structure of catchment areas, and instead aims to produce partitions that reflect stable, naturally occurring relationships within pre-existing patterns of healthcare usage. As expected, the coarser partitions, which naturally correspond to multi-hospital catchment areas, do increase coverage as observed in our results. However, the advantages of such coverage need to be balanced by the costs associated with increased concentration and centralisation. Such costs need to be accounted for based on different operational priorities by managers and policy makers. The method thus provides a tool to explore the relative benefits of information-sharing at different levels of coarsening in a data-informed manner based on patterns of hospitalisation usage, rather than geographical or administrative divisions. The resulting catchment areas must therefore be appraised in a wider policy context, in which they would be implemented, reflecting local organisational challenges and priorities that may favour one proposed configuration over another.

As populations age and the complexity of medical care increases, it becomes imperative for secondary care providers to reach out into the communities they serve.^16^ Catchment areas derived from multiscale community detection facilitate parsimonious collaboration between hospitals to provide comprehensive, meaningful interventions in the communities they serve. Their ability to produce meaningful representations of how secondary care organisations may collaborate with one another to reach out into their local communities may serve as an important tool as the National Health Service in England seeks to formalise the local accountability of hospitals.

Multiscale community detection is a novel, data-driven, unsupervised method to generate mutually exclusive, collectively exhaustive catchment areas based on patterns of patient presentation. Its application to data on planned orthopaedic care in England shows an improvement in the identification of representative catchment areas over existing methods. The advantage of this method is particularly in permitting multiple hospitals to collectively share a single defined catchment area. In doing so, this offers the opportunity for dilute health care markets in urban areas to be comprehensively, parsimoniously covered by collaborations between institutions. As the National Health Service in England seeks to enhance collaboration between hospital and community care, this methodology of Markov Stability multiscale community detection offers a powerful tool to understand how such collaborations should be structured.

## Acknowledgements

This article is independent research supported by grants from The Peter Sowerby Foundation and the National Institute for Health Research (NIHR) Imperial Patient Safety and Translational Research Centre (PSTRC). Infrastructure support for this work was provided by the NIHR Imperial Biomedical Research Centre (BRC). MB acknowledges support from EPSRC grant EP/N014529/1 supporting the EPSRC Centre for Mathematics of Precision Healthcare. The views expressed in this publication are those of the author(s) and not necessarily those of the NHS, the National Institute for Health Research or the Department of Health. The authors declare no competing interests in regard to this study.

## Author Contributions

JC was involved in all aspects of the study. MB and AD were involved in the planning, interpretation, writing and reviewing of the study.

